# Impact of chronic transcranial Random-Noise Stimulation (tRNS) on prefrontal cortex excitation-inhibition balance in juvenile mice

**DOI:** 10.1101/2020.09.04.282889

**Authors:** Carlos A. Sánchez-León, Álvaro Sánchez-López, María A. Gómez-Climent, Isabel Cordones, Roi Cohen Kadosh, Javier Márquez-Ruiz

## Abstract

Transcranial random noise stimulation (tRNS), a non-invasive neuromodulatory technique capable of altering cortical activity, has been proposed to improve the signal-to-noise ratio at the neuronal level and the sensitivity of the neurons following an inverted U-function. The aim of this study was to examine the effects of tRNS on vGLUT1 and GAD 65-67 and its safety in terms of pathological changes. For that, juvenile mice were randomly distributed in three different groups: “tRNS 1x” receiving tRNS at the density current used in humans (0.3 A/m^2^, 20 min), “tRNS 100x” receiving tRNS at two orders of magnitude higher (30.0 A/m^2^, 20 min) and “sham” (0.3 A/m^2^, 15 s). Nine tRNS sessions during five weeks were administered to the prefrontal cortex of alert animals. No detectable tissue macroscopic lesions were observed after tRNS sessions. Post-stimulation immunohistochemical analysis of GAD 65-67 and vGLUT1 immunoreactivity showed a reduced GAD 65-67 immunoreactivity levels in the region directly beneath the electrode for tRNS 1x group with no significant effects in the tRNS 100x nor sham group. The observed results points to an excitatory effect associated with a decrease in GABA levels in absence of major histopathological alterations providing a novel mechanistic explanation for tRNS effects.

## Introduction

Transcranial random noise stimulation (tRNS) delivers a painless, weak current at random, constantly changing frequencies usually between 101Hz and 640Hz. Previous research has highlighted the benefit of random noise and in particular it has shown that tRNS can alter cortical activity during (Snowball et al., 2013) and after stimulation (Terney et al., 2008), with some behavioural and neural effects lasting up to several months (Cappelletti et al., 2013; Snowball et al., 2013; Pasqualotto, 2016; Frank et al., 2018; Brevet-Aeby et al., 2019; Herpich et al., 2019).

Compared to more familiar methods, such as transcranial magnetic stimulation, transcranial direct current stimulation (tDCS) or transcranial alternating current stimulation (tACS), tRNS is considered the most comfortable intervention technique for participants, which is a key advantage for use with cognitive training and effective blinding (i.e., whether the participant receives sham or active tRNS). For example, the 50% perception threshold for tDCS was set at 0.4 mA, and at 1.2 mA in the case of tRNS (Ambrus et al., 2010). In addition, tRNS exhibited long-lasting effects in several studies (Cappelletti et al., 2013; Snowball et al., 2013; Pasqualotto, 2016; Herpich et al., 2019; Brevet-Aeby et al, 2019; Frank et al., 2018). tRNS is also polarity-independent, with both electrodes (Terney et al., 2008), or at least one (Snowball et al., 2013) inducing excitatory effects (when the current is set to 1mA). tRNS is less sensitive to cortical folding than other neurostimulation methods (Terney et al., 2008), reducing the impact of anatomical variations between participants.

One of the suggested mechanisms in tRNS is stochastic resonance (Terney et al., 2008; van der Groen and Wenderoth, 2016; Fertonani and Miniussi, 2017; Harty and Cohen Kadosh, 2019). According to this framework in non-ideal and non-linear systems, such as the brain, noise can be beneficial. This fact has been shown in a number of research fields, including perception, ecology, and engineering (McDonnell and Abbott, 2009; McDonnell and Ward, 2011). In all of these cases, when noise is applied to a subthreshold signal/input it will improve performance/output. Critically, the noise needs to be at a specific level to yield optimal gain as the addition of too much noise can be non-beneficial. The main assumption is that tRNS induces noise in the neural system and as a consequence will improve the signal-to-noise ratio at the neuronal level and the sensitivity of the neurons (Fertonani et al., 2011). This idea is based on the most prevalent hypothesis of noise-induced improvement in multiple disciplines (McDonnell and Abbott, 2009). In the present experiment, the prediction is that random noise-based neurostimulation should influence the brain excitability non-linearly. That is, the changes in the tRNS parameters should follow an inverted U-function in that the output metric is very small for high and no noise (sham stimulation), while low noise level provides more optimal output (McDonnell and Abbott, 2009).

It is surprising that despite the promising results of tRNS in human-based research, in some cases even more than more popular methods such as tDCS and tACS (Fertonani et al., 2011; Brem et al., 2018; Simonsmeier et al., 2018; Berger et al., 2019) its mechanisms are relatively unclear, and are scarce compared to other methods such as tDCS and tACS. In addition, its safety based on animal-based research is lacking. This is a point of potential concern as tRNS is now being used in the case of neurodevelopmental disorders (Looi et al., 2017; Berger et al., 2019). Therefore, the motivation in this study was to mirror a promising protocol of tRNS that was used during a cognitive training in atypical developing children, in order to examine its effects at vGLUT1 and GAD 65-67 that are involved in neuroplasticity and its safety in terms of pathological changes induced by tRNS.

To do so we used the protocol published by Looi et al (2017), who administrated chronic tRNS in the shape of 9 tRNS sessions during five weeks to children with mathematical learning difficulties. Juvenile mice were randomly distributed in three different groups: 1) “tRNS 1x” group: receiving tRNS at the same density current as in Looi et al., (2017) (0.3 A/m^2^, 20 min, n = 5), 2) “tRNS 100x” group: receiving tRNS at two order of magnitude higher than those commonly used in human tRNS experiments (30.0 A/m^2^, 20 min, n = 6), and 3) “sham” group: receiving the same tRNS than tRNS 1x group except for time duration that was 15 s instead of 20 min (n = 5).

## Methods

### Animals

Experiment was carried out on 6 weeks old males C57 mice (University of Seville, Spain) weighing 28–35 g. Before and after surgery, the animals were kept in the same room but placed in independent cages. The animals were maintained on a 12-h light/12-h dark cycle with continuously controlled humidity (55 ± 5%) and temperature (21 ± 1 °C). All experimental procedures were carried out in accordance with European Union guidelines (2010/63/CE) and following Spanish regulations (RD 53/2013) for the use of laboratory animals in chronic experiments. In addition, these experiments were submitted to and approved by the local Ethics Committee of the Pablo de Olavide University (Seville, Spain).

### Surgery

Animals were anesthetized with a ketamine–xylazine mixture (Ketaset, 100 mg/ml, Zoetis, NJ., USA; Rompun, 20 mg/ml, Bayer, Leverkusen, Germany) at an initial dosage of 0.1 ml/20 g. Under aseptic conditions, an anteroposterior (AP) incision in the skin along the midline of the head, from the front leading edge to the lambdoid suture, was performed. Subsequently, the periosteum of the exposed surface of the skull was removed and the bone was washed with sterile saline. The animal’s head was correctly positioned in an stereotaxic frame (David Kopf Instruments, CA, USA) to mark the position of bregma as stereotaxic zero. For tRNS administration, a polyethylene tubing (3 mm inner diameter; A-M Systems), which acted as the active electrode for tRNS, was placed over the skull centered on the prefrontal cortex (AP = + 1.8 mm; Lateral = 0 mm; relative to bregma (Paxinos and Franklin, 2004)) (Fig. 1A). Once placed in the correct coordinates, the tube was externally covered with dental cement (DuraLay, Ill., USA). Finally, a head-holding system was implanted, consisting of three bolts screwed to the skull and a bolt placed over the skull upside down and perpendicular to the horizontal plane to allow for head fixation during the tRNS protocol. The complete holding system was cemented to the skull.

**Figure 1.**
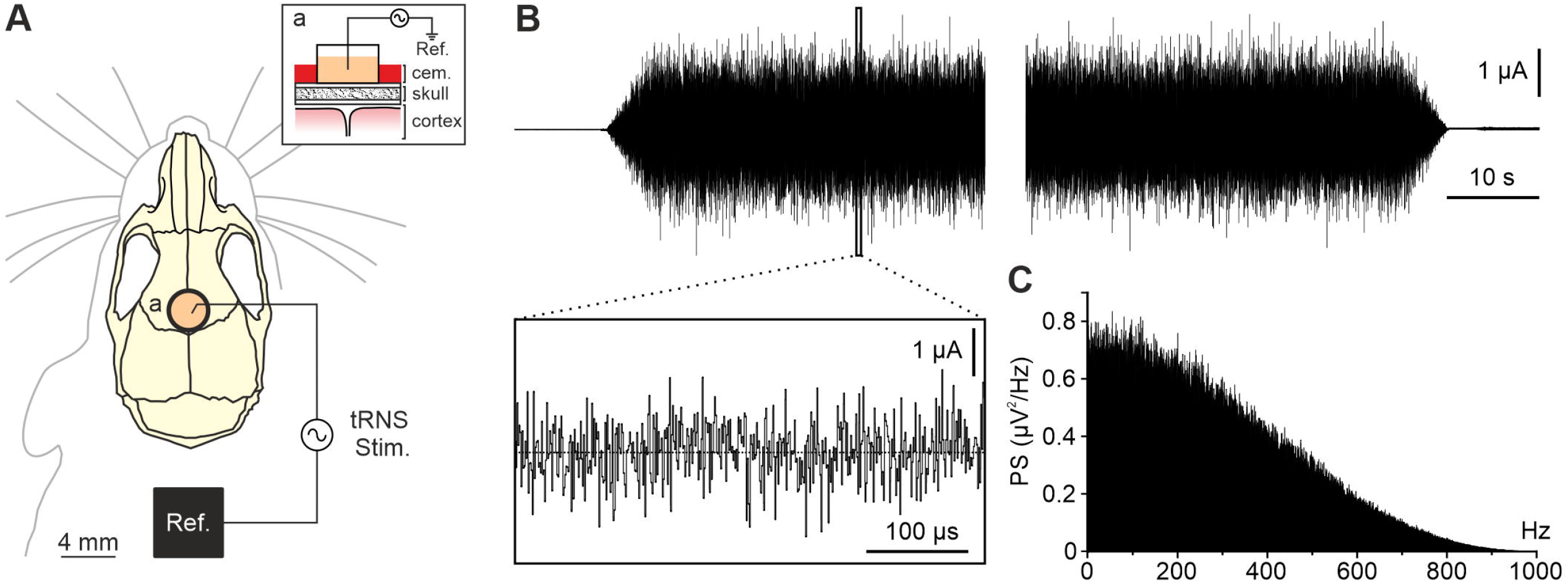
Experimental preparation and tRNS protocol. A) Illustration of the experimental design showing active electrode (a) and reference (Ref.) locations for tRNS (tRNS Stim.) in the PFC of alert mice. Inset (a) shows a sagittal schematic view of the stimulating site. B) The figure shows the current waveform passing through active and reference electrodes during tRNS protocol. Five seconds ramp up (upper trace, left) and ramp down (upper trace, right) were used at the beginning and at the end of the 20 min stimulation. Inset box shows an expanded (x-axis) view of the waveform. C) Power spectrum of the recorded stimulation signal.

### tRNS sessions

Stimulating sessions began at least one week after surgery. The animals were placed over a treadmill to reduce the stress in head-fixed condition and the head was fixed to the recording table by means of the implanted head-holding system. The different protocols for tRNS were designed in StarStim tES-EEG system (Neuroelectrics, Barcelona, Spain) and sent to a battery-driven linear stimulus isolator (WPI A395, Fl, USA). The tubing used as active electrode was filled with electrogel (Electro-Cap International, OH., USA) and a metallic electrode from the stimulus isolator was immersed in it. tRNS was applied between the active electrode over PFC and a reference electrode consisting on a rubber rectangle (6 cm^2^) attached to the back of the mouse and moisten with electrogel. To characterize long-lasting histological changes, tRNS was delivered during 20 min (including 5 s ramp-up and 5 s ramp-down, 0.1–800Hz) at 0.3 A/m^2^ in the first group of animals (n = 5), 30 A/m^2^ in the second group (n = 6) and for 15 seconds (including 5 s ramp-up and 5 s rampdown) at 0.3 A/m^2^ in the sham group (n = 5). Mice were stimulated twice a week receiving a total of 9 sessions.

### Histology

To characterize potential histological changes on PFC glutamate and GABA levels after tRNS at different current densities mice were deeply anesthetized with ketamine–xylazine mixture (Ketaset, 100 mg/ml; Rompun, 20 mg/ml) 15 min after last tRNS session and perfused transcardially with 0.9% saline followed by 4% paraformaldehyde (PanReac, Barcelona, Spain) in phosphate buffer pH 7.4 (PB). The brains were removed and stored at 4 °C in the 4% paraformaldehyde solution for 24 hours, cryoprotected in 30% sucrose in PBS the next 48 hours, and then cut into 50 μm coronal slices with a freezing microtome (CM1520, Leica, Wetzlar, Germany). Slices underwent a toluidine blue or immunohistochemical staining protocol. Sections were processed “free-floating” and passed through all procedures simultaneously to minimize differences in immunohistochemical staining. After three washes of 10 min with PBS, sections were blocked with 10% Normal Donkey Serum (NDS, 566460, Merck, Darmstadt, Germany) in PBS with 0.2% Triton X-100 (Sigma-Aldrich, Mo., USA) (PBS-Tx-10% NDS) and then incubated overnight at room temperature in darkness with the primary antibody solution containing mouse anti-vesicular Glutamate Transporter 1 (vGlut1, 1:1000, MAB5502, Merck) and rabbit anti-Glutamate Decarboxylase 65-67 (GAD 65-67, 1:1000, AB1511, Merck). After three washes, sections were incubated for 1 hour at room temperature in darkness with appropriate secondary antibodies: Alexa Fluor 488 donkey anti-mouse IgG (H+L) (1:400, A21202, Thermo Fisher Scientific, Mass., USA), Alexa Fluor 555 donkey anti-rabbit IgG (H+L) (1:400, A31572, Thermo Fisher Scientific) in PBS-Tx-5% NDS. After three washes with PBS, sections were stained with Hoechst 33342 dye (Merck Millipore, Billerica, MA, USA) and mounted on glass slides, and coverslipped using Dako Fluorescence Mounting Medium (Dako North America, CA., USA). For confocal imaging, an *in vivo* confocal microscope (A1R HD25, Nikon, Tokyo, Japan) was used. Z-series of optical sections (0.5 μm apart) were obtained using the sequential scanning mode.

### Data analysis

Confocal images were processed in ImageJ (https://imagej.nih.gov/ij/) with the image-processing package Fiji (http://fiji.sc/Fiji) using a custom-built macro. To subtract fluorescence background noise five square regions of interest (ROI) of 30 x 30 pixels (26.22 μm^2^) were placed over unlabeled nuclei in each image, and the obtained maximum brightness average was set as the minimum value for “setThreshold”, so the pixels with values lower than the average were considered as non-fluorescent. Then, a copy of the image was converted to binary to visually validate the procedure, and the complete process was repeated until the threshold properly discriminated our signal from the noise. To analyze particles, five square regions of interest (ROI) of 100 x 100 pixels (291.31 μm^2^) were randomly placed over regions absent of nuclei or unspecific noise (as for example blood vessels). Each image inside the ROI was converted to binary and the “Analyze Particles” command was used to count and measure aggregates of vGlut1 and GAD65-67. Particles were sorted as small (size = 10-25), medium (size = 26-46) or big (size = 47-100) and each category was averaged across the five ROI to obtain one value per hemisphere per animal.

### Statistical analysis

To examine stimulation effects, we used linear mixed effects models, which account for within-subject correlations more optimally compared to ANOVA and automatically handle missing values, allowing maximum use of available data (Seltman, 2009). We used the R-package nlme (Pinheiro et al., 2017) to perform the linear mixed effects analysis with maximized log-likelihood on the outcome measures. We examined outcomes (GAD 65-67, VGLUT1) as a function of stimulation condition, regions, and their interaction as predictors. We used sham and Area 4 (which was the deeper region, which is assumed to be least impacted by stimulation, if at all) as the reference variable. For all the measures we verified that the residuals were normally distributed using a q-q plot and the Shapiro–Wilk normality test. The results are shown as mean ± SEM. Statistical significance was set at p < 0.05 in all cases.

### Data availability

Data is available upon reasonable request from the corresponding author.

## Results

To explore for potential changes in the excitation/inhibition balance in PFC after exposition to tRNS, young mice (6 weeks old) were prepared for chronic stimulation in alert head restrained condition (Fig. 1A). The three first days after surgery recovery were used to habituate the mice to the treadmill and head-fixed condition. During tRNS sessions, the animals were placed over a treadmill and the head fixed to the recording table by means of the implanted head-holding system. The plastic tube chronically implanted on the skull over PFC was cleaned and filled with electrogel. A chlorinated-silver wire was inserted in the tube and a square rubber electrode (6 cm^2^) moisten with electrogel was attached to the back of the animal and used as reference electrode. After tRNS session, silver wire and rubber electrode were removed, and the animal placed again in its cage.

After 9^th^ tRNS session, mice from each group were transcardially perfused 15 min after the end of the protocol and the brains were processed for toluidine blue and immunohistological analysis. A first histological evaluation was carried out in the toluidine blue stained slices with light microscopy in order to find pathological changes such as oedema, necrosis and haematoma and for cellular alterations (Liebetanz et al., 2009, Jackson et al., 2017). The figure 2 shows a low-magnification photomicrograph coronal reconstruction at the PFC level for three representative animals from sham (Fig. 2A), tRNS 1x (Fig. 2B) and tRNS 100x (Fig. 2C) groups. No signs of tRNS-induced neurotrauma was observed for any of the applied current densities (0.3 A/m^2^ and 30.0 A/m^2^) nor sham condition. No histopathological alterations were observed at higher magnification in any of the four selected areas from different animal groups (1-4 insets in Fig. 2A-C).

**Figure 2.**
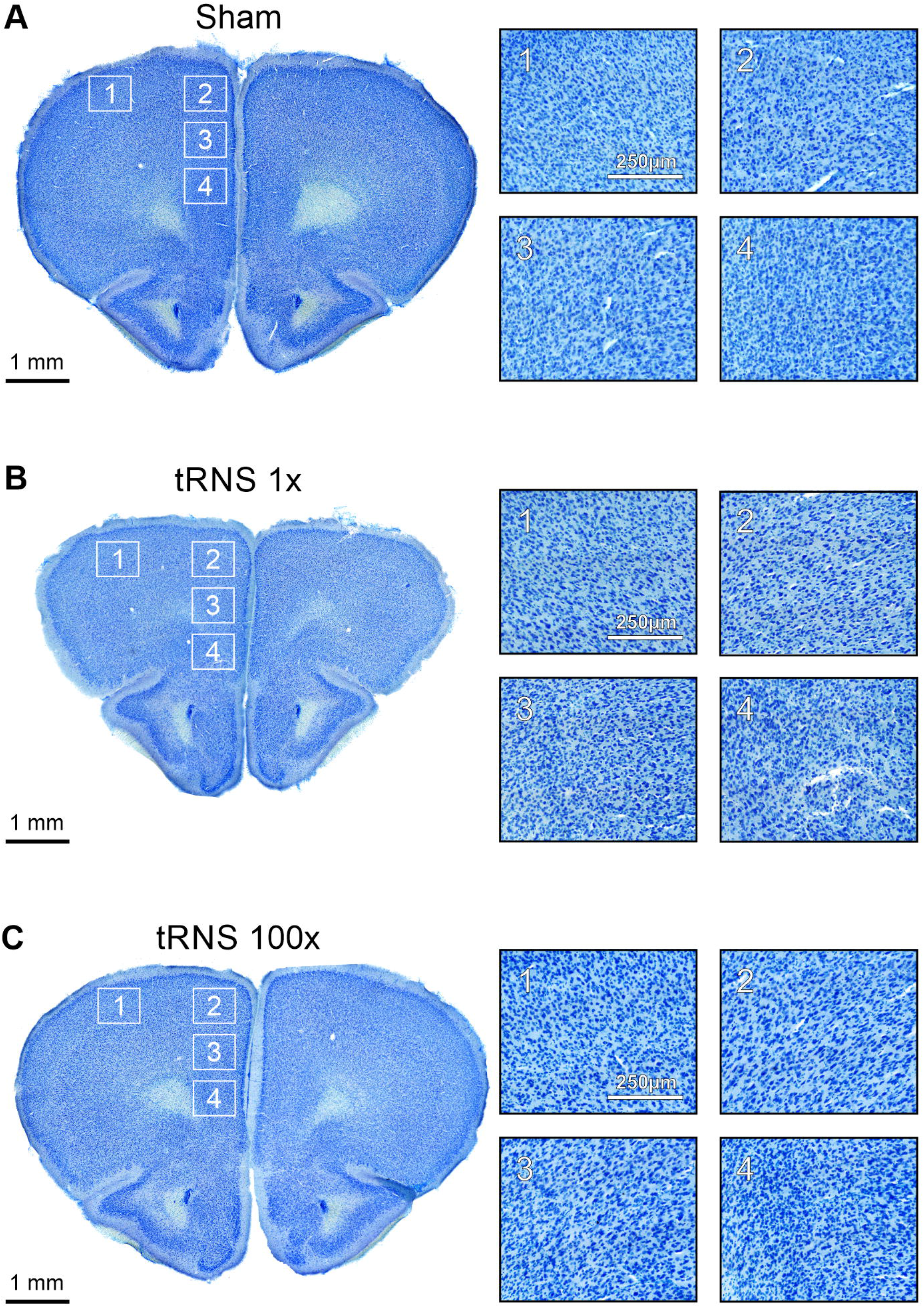
Histological analysis of the stimulated cortical regions. Low-magnification photomicrograph of a coronal section at PFC level, stained with toluidine blue, for three representative animals from sham (A), tRNS 1x (B) and tRNS 100x. Right column shows higher-magnification microphotographs corresponding to the four squared areas represented at left (1-4). No histopathological alterations were found in any of the stimulated groups.

In order to explore the potential impact of tRNS on cortical markers of excitation and inhibition we performed a post-stimulation immunohistochemical analysis of GAD 65-67 and vGLUT1 immunoreactivity in the stimulated PFC region. The number of GAD 65-67 and vGLUT1 positive clusters of puncta in four different areas of the stimulated PFC were analyzed in the sham, tRNS 1x and tRNS 100x groups and compared (Fig. 3A). For GAD 65-67, we found a significant interaction between tRNS 1x and Area 1 and Area 2 (tRNS 1x*Area 1: B = −10.08, SE = 4.29, t(39) = −2.35, p = 0.024; tRNS 1x*Area 2: B = −17.16, SE = 4.29, t(39) = −4, p = 0.0003, Table 1). The results as plotted in figure 3B show that compared to sham group for tRNS 1x there was a reduced GAD 65-67 immunoreactivity levels in the region directly beneath the electrode and on the surface nearby (1 and 2 areas in Fig. 3A). In contrast, there were no significant effects in a deeper region, or when tRNS was increased (tRNS 100x). For vGLUT1, none of the effects were significant (Fig. 3C, Table 2). When we applied corrections for multiple comparisons, the interaction between tRNS 1x and Area 2 was still significant (p < 0.0066, Benjamini-Hochberg correction with a false discovery rate=0.05). Figure 3 shows representative confocal images from the “Area 2” in the sham, tRNS 1x and tRNS 100x groups for GAD 65-67 (Fig. 3D-F) and vGLUT1 (Fig. 3G-I).

**Figure 3.**
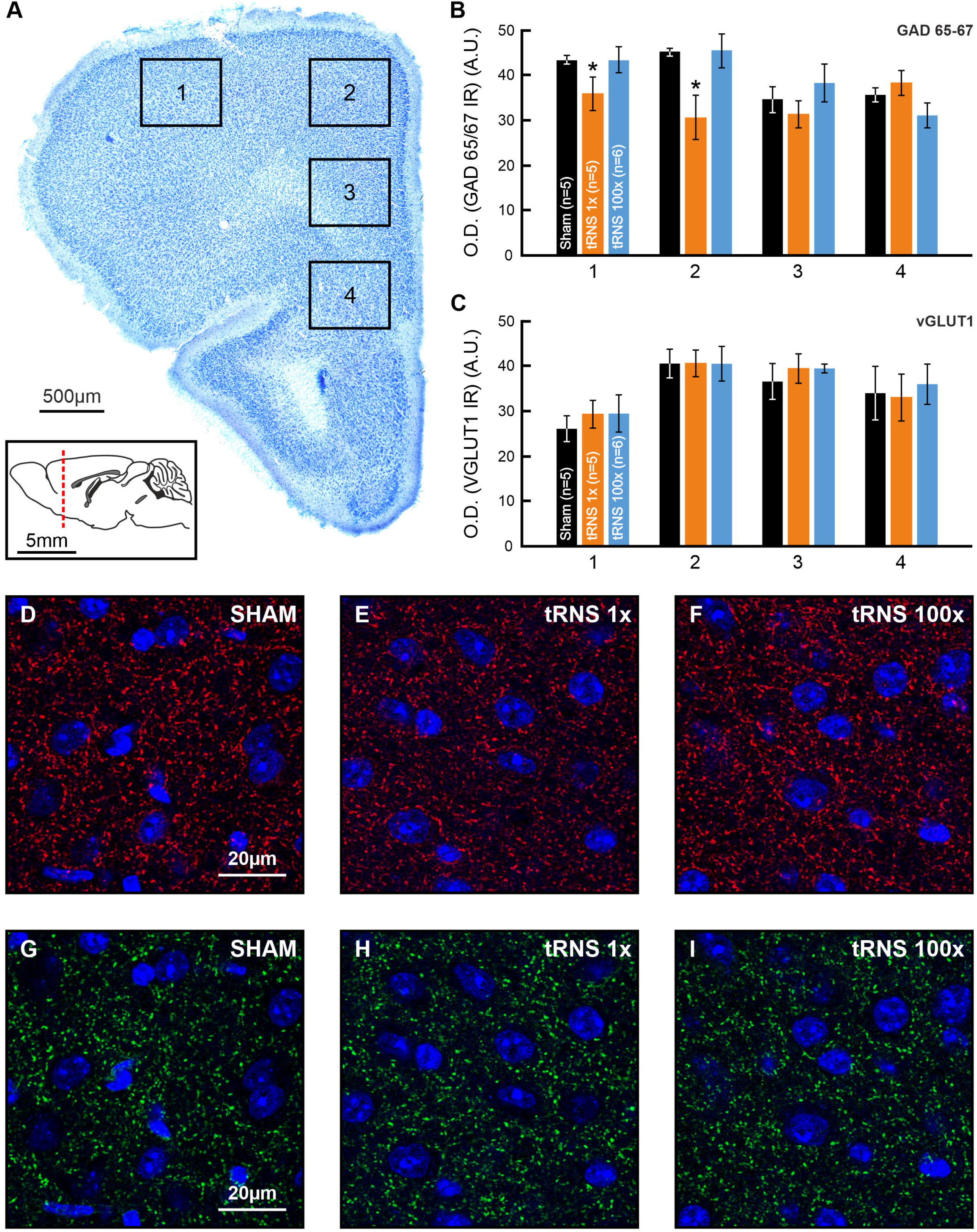
Immunohistochemical changes after chronic tRNS. A) Low-magnification photomicrograph of a coronal section at PFC level, representing the four immunohistochemically analyzed areas (1-4). Sagittal mouse brain scheme indicates the level from which the analyzed slices were obtained. B,C) Quantification and statistics (bar charts) of GAD 65-67 (B) or vGLUT1 immunoreactivity (C) in the four analyzed areas (1-4, indicated in A) for sham (n = 5), tRNS 1x (n = 5) and tRNS 100x (n = 6) groups after 9 tRNS sessions (20 min). D-I) Confocal photomicrographs of GAD 65-67 (D-F) and vGLUT1 (G-I) immunoreactivity and Hoechst fluorescence in “area 2” of PFC for representative animals from sham, tRNS 1x and tRNS 100x groups. Error bars represent SEM. GAD 65-67: Glutamic acid decarboxylase isoforms 65 and 67; vGLUT1: vesicular glutamate transporter 1; OD: optical density; IR: immunoreactivity; A.U.: arbitrary units.

**Table 1.** Beta Weights of the Regression Model with GAD 65-67 values as the Outcome Measure, with sham as the reference variable and Area 4 as the reference region. The results indicate a significant effect for tRNS 1x with Area due to greater reduction GAD 65-67 values for tRNS 1x in comparison to Sham in Areas 1 and 2. SE=standard errors, DF=degrees of freedom.

|  | Beta | SE | DF | t-value | p-value |
| --- | --- | --- | --- | --- | --- |
| (Intercept) | 35.68 | 3.28 | 39 | 10.88 | <0.0001 |
| tRNS 1x | 2.56 | 4.64 | 13 | 0.55 | 0.5903 |
| tRNS 100x | -4.58 | 4.44 | 13 | -1.03 | 0.3211 |
| Area1 | 7.76 | 3.03 | 39 | 2.56 | 0.0146 |
| Area2 | 9.36 | 3.03 | 39 | 3.08 | 0.0037 |
| Area3 | -1.04 | 3.03 | 39 | -0.34 | 0.7337 |
| tRNS 1x*Area1 | -10.08 | 4.29 | 39 | -2.35 | 0.024 |
| tRNS 100x:Area1 | 4.54 | 4.11 | 39 | 1.10 | 0.276 |
| tRNS 1x*Area2 | -17.16 | 4.29 | 39 | -4.00 | 0.0003 |
| tRNS 100x:Area2 | 4.87 | 4.11 | 39 | 1.19 | 0.2428 |
| tRNS 1x*Area3 | -5.92 | 4.29 | 39 | -1.38 | 0.1756 |
| tRNS 100x:Area3 | 8.31 | 4.11 | 39 | 2.02 | 0.0501 |

**Table 2.** Beta Weights of the Regression Model with vGLUT1 values as the Outcome Measure, with sham as the reference variable and Area 1 as the reference region. SE=standard errors, DF=degrees of freedom.

|  | Beta | SE | DF | t-value | p-value |
| --- | --- | --- | --- | --- | --- |
| (Intercept) | 33.96 | 3.98 | 39 | 8.53 | <0.0001 |
| tRNS 1x | -0.96 | 5.63 | 13 | -0.17 | 0.8672 |
| tRNS 100x | 2.01 | 5.39 | 13 | 0.37 | 0.7157 |
| Area1 | -7.92 | 5.26 | 39 | -1.51 | 0.1402 |
| Area2 | 6.6 | 5.26 | 39 | 1.25 | 0.2171 |
| Area3 | 2.6 | 5.26 | 39 | 0.49 | 0.6239 |
| tRNS 1x*Area1 | 4.2 | 7.44 | 39 | 0.56 | 0.5756 |
| tRNS 100x:Area1 | 1.39 | 7.12 | 39 | 0.19 | 0.8466 |
| tRNS 1x*Area2 | 1 | 7.44 | 39 | 0.13 | 0.8938 |
| tRNS 100x*Area2 | -2.03 | 7.12 | 39 | -0.29 | 0.7768 |
| tRNS 1x*Area3 | 3.84 | 7.44 | 39 | 0.51 | 0.6086 |
| tRNS 100x*Area3 | 0.9 | 7.12 | 39 | 0.13 | 0.9001 |

## Discussion

The present histological results in young mice suggest that tRNS applied at low-density currents is capable of increasing excitability by decreasing GABA levels in a focalized way. These results could be of crucial importance for human tRNS studies suggesting that a decrease in GABA levels could be mediating the behavioral enhancement observed in previous studies (Terney et al., 2008; Fertonani et al., 2011; Cappelletti et al., 2013; Snowball et al., 2013; Pasqualotto, 2016; van der Groen and Wenderoth, 2016; Looi et al., 2017; Brem et al., 2018; Evans et al., 2018; Frank et al., 2018; Brevet-Aeby et al, 2019; Herpich et al., 2019; Berger et al., 2019; Harty & Cohen Kadosh et al., 2019; Sheffield et al., 2020) and that the technique do not produce major histopathological alterations at the density currents used in the study.

The first aim of this study was to evaluate potential pathological changes after chronic exposition to tRNS in young animals. No histopathological alterations were observed for any of the applied current densities (0.3 A/m^2^ and 30.0 A/m^2^). Previous safety limits studies in rodents estimated no detectable tissue macroscopic lesions below a density current of 28.6 A/m^2^ for cathodal (Liebetanz et al., 2009) and 20 A/m^2^ for anodal tDCS (Jackson et al., 2017) with brain tissue remaining lesion free when charge density (density current x time) was set below 52,400 C/m^2^ and 72,000 C/m^2^, respectively. In the present work, we used 339.5 C/m^2^ for tRNS 1x and 33,953.1 C/m^2^ tRNS 100x group, supporting above mentioned safety limits studies (Liebetanz et al., 2009; Jackson et al., 2017). Interestingly, Liebetanz and colleagues (2009) also report that chronic tDCS application at 68,571 C/m^2^ for 5 consecutive days produced no tissue damage. We obtained similar results in this study with no macroscopic lesions after 9 tRNS sessions. Nevertheless, beyond macroscopic lesions, tDCS has been demonstrated to change different molecular mediators involved in immune- and inflammatory processes in rats (Rueger et al., 2012; Pikhovych et al., 2016; Rabenstein et al., 2019) even when no apparent cortical lesions were observed. It is still unknown how tRNS could affects these neuroinflammatory processes, and we hope to extend this knowledge in future studies.

Our second aim was to explore the impact of chronic tRNS on excitation-inhibition balance through glutamate- and GABA-related markers. tRNS has been proposed to increase the excitability of brain cortex during and after stimulation (Terney et al., 2008; Snowball et al., 2013; Pasqualotto, 2016; Frank et al., 2018; Herpich et al., 2019; Brevet-Aeby et al, 2019). Nevertheless, brain mechanisms remain unclear and animal model experiments are missing in the literature (Antal and Herrmann, 2016). While on-line effects have been related with the potentiation of voltage-gated Na^+^ channels (Schoen and Fromherz, 2008; Terney et al., 2008; Chaieb et al., 2015) and stochastic resonance (Terney et al., 2008; van der Groen and Wenderoth, 2016; Fertonani and Miniussi, 2017), brain mechanisms underlying tRNS long-term effects are still missed. We show here that chronic tRNS over PFC decreased GAD 65-67 immunoreactivity related with GABA levels at low density current (0.3 A/m^2^), similar to those used in humans, with no effects for sham nor high density current stimulation (30.0 A/m^2^). This reduction in the inhibitory neurotransmitter could lead to an increase in the cortical excitability as already observed in humans after anodal tDCS in the primary motor cortex (Stagg et al., 2009; Bachtiar et al., 2018; Patel et al., 2019). Interestingly, GABA_A_ agonist lorazepam has been shown to suppress tRNS-induced cortical excitability increases in human subjects whereas NMDA receptor agonist D-cycloserine, the NMDA receptor antagonist dextromethorphan and the D2/D3 receptor agonist ropinirole have no significant effects (Chaieb et al., 2015). The results shown in the present study were dependent on the tRNS applied density current, being evident for low density currents but not statistically significant for high-density current or sham stimulation. This non-linear neurostimulation effects fits well with the proposed stochastic resonance predicting and optimal output for low noise levels following an inverted U-function (McDonnell and Abbott, 2009), and matched the enhancement effect found by tRNS in previous human-based experiments (van der Groen and Wenderoth, 2016; Harty and Cohen Kadosh, 2019). On the other hand, statistically significant effects were restricted to the PFC area just under the active electrode (Area 1 and 2 in Fig. 3A) presenting a higher GAD 65-67 reduction in the Area 2 placed near from the electrode center. Considering the anatomical characteristics of the mouse PFC, significant effects were restricted to medial prefrontal cortex, and more specifically to anterior cingulate cortex, with prelimbic and infralimbic cortices not showing significant GAD 65-67 reduction (Dalley et al., 2004; Bicks et al., 2015). Absence of long-term histological effects in Area 3 and 4 could be due to the focality of tRNS, with electric fields generated behaving in a linear ohmic manner, reaching higher intensity values close to the active electrode position (Sánchez-León et al., 2020). Nevertheless, we cannot discard a potential impact of neuronal axo-dendritic orientation (with respect to the active electrode position) in the final observed results (Bikson et al., 2004; Radman et al., 2009; Kabakov et al., 2012; Rahman et al., 2013), with anterior cingulate cortex neurons differently oriented than deeper prefrontal cortex regions. The axo-dendritic angle of cingulate cortex neurons could confer to this neuronal population a higher sensitivity to the electric fields than prelimbic and infralimbic neurons or even predominantly affects to different neuronal types (e.g., pyramidal, basket, chandelier cells) in each region. However, it is important to note that it was suggested that tRNS is less sensitive to cortical folding than other neurostimulation methods (Terney et al., 2008).

The present results, pointing to an excitatory effect associated with a decrease in GABA levels, constitute a first step toward the understanding of basic mechanisms of tRNS in alert animals. Nevertheless, future experiments combining tRNS with electrophysiological recordings and behavioral performance are urgently needed. A recent tRNS-electroencephalography study in accordance with our results, show that the tRNS effect on human arithmetic learning depends on the individual’s excitation/inhibition levels during rest and task performance (Sheffield et al., 2020). Notably, those with lower excitation/inhibition levels benefited more from the potential excitatory effect induced by tRNS (Sheffield et al., 2020). However, that study did not find that a single-session of tRNS alters the excitation/inhibition balance, and future studies could examine if multiple sessions are required for such effect.

*Adolescence* and *Juvenile* terms have been used in rodents to cover the whole time span from weaning at postnatal day 21 to adulthood at postnatal day 60 (Babikian et al., 2010). Taken into account the age of the participating animals at the beginning of the experiment (~6 weeks old) and the duration of the tRNS protocol (4.5 weeks) we chronically stimulated from postnatal day 42 to postnatal day 74. The time interval covered during the applied tRNS protocol has been compared with 12-18 years old in humans (where a synapse density reduction, cognitive-dependent circuitry refinement and white matter volume increase have been described) and adulthood (> 20 years old, where neurotransmitter and synapse density reaches adult levels) (see Semple et al., 2013 for a review). As a result, the stimulated time window was delayed with respect to Looi and colleagues (2017) where 9 tRNS sessions were applied to 8-10 years old children for five weeks. While the results applied to older participants than those in Looi et al. (2017), the present results provide unique and novel information on the impact of tRNS on the developing brain. We assume that the mechanisms of the tRNS effect we observed will be similar in younger and older participants. However, further data would be necessary to examine our view.

Animal models have been successfully used to disentangle tES mechanisms in the past (see Sánchez-León et al., 2018 for a review). Differently from other popular animal models in the tES field like rats, mice offer the opportunity to use transgenic animals for dissecting tES effects at the neuronal population level or to work with murine models resembling human pathologies. Thus, alert mice models have been successfully used to figuring out tES mechanisms offering a unique opportunity to deepen in neuronal (Sun et al., 2020; Sánchez-León et al., 2020) and glial (Monai et al., 2016; Mishima et al., 2019) response to tES, to study new electrical stimulation paradigms (Grossman et al., 2017), or to explore its potential therapeutic application in different pathologies (Lu et al., 2015; Souza et al., 2018; Peanlikhit et al., 2017). Shedding light on neuronal mechanisms governing tRNS effects in animals will be crucial to understand the impact of this non-invasive technique in the human brain and to optimize stimulation protocols in a comprehensive way.

